# Amyloid beta aggregation promoted by iron leads to neuronal loss in an *ex vivo* model of Alzheimer’s disease

**DOI:** 10.1101/2025.07.11.664035

**Authors:** Sowmya Sunkara, Rahul Krishnamoorthy, Snježana Radulović, Marlene Leoni, Silke Patz, Walter Goessler, Walter A Kaufmann, Anna Birkl-Toeglhofer, Johannes Haybaeck, Gerd Leitinger

## Abstract

Alzheimer’s disease (AD) is a progressive neurodegenerative disorder characterized by amyloid beta (Aβ) plaques and neurofibrillary tangles. Despite well-established iron accumulation in the AD brain, its role in exacerbating Aβ toxicity is often overlooked in therapeutic research. We developed a 3D *ex vivo* organotypic brain slice cultures (OBSC) with Aβ monomers and ferric citrate to mimic Aβ deposits and iron overload to investigate the impact of excess iron on Aβ toxicity in pig and human brains. Light and electron microscopy, biochemical assays, and multiple regression modeling were employed to assess iron-mediated Aβ toxicity in neurons and glial cells.

We show that OBSC offer a close approximation of *in vivo* morphological and physiological properties and can retain both neurons and glial cells for extended periods, and respond to experimental manipulations. We show that iron promotes Aβ fibrillization into long fibrils, with this process further influenced by temperature. Aβ selectively accumulated in neurons, leading to their death, sparing glial cells. In contrast, Iron, though generally toxic to neurons, exhibited unspecific cytotoxicity. Notably, the combined presence of Aβ and iron synergistically increased neuronal death while reducing glial cell loss. Correlation analysis revealed that this synergic interaction enhances the toxicity of each other in a mutual fashion – Aβ directs the neuronal toxicity while iron promotes Aβ fibrillization, leading to targeted neuronal loss.

In conclusion, our findings emphasize the critical role of excess iron and Aβ in driving neuronal death in AD, underlining the importance of targeting iron accumulation along with Aβ clearance but also addressing in future AD therapies, while also supporting our OBSC model as a valuable platform for studying the same.

## Introduction

Alzheimer’s disease (AD) is a progressive neurodegenerative disorder that gradually disrupts the connections between brain cells potentially affecting brain functioning. AD is characterized by two main biomarkers - amyloid beta (Aβ) deposits, and tau neurofibrillary tangles. Both are already detectable in the preclinical stage, when neuroinflammation and neuronal loss start to manifest themselves (1,2). Of these, Aβ has long been regarded as a crucial biomarker since Aβ is the first to be detected in the course of the disease (71,92) and triggers neurodegeneration in the AD brain (71).

Aβ is derived via cleavage from amyloid precursor protein (APP), exists in two primary isoforms: Aβ_1-40_ and Aβ_1-42_ (5,6). Among these, Aβ_1-42_ is the most toxic and closely associated with AD pathology (7–10). These Aβ monomers assemble into various structures, ranging from five nanometres to several microns in size depending on the type of species formed. Aβ forms globular oligomers, resembling small beads with an average diameter of 10 nm (1) or organize into small linear to curved forms, typically measuring 40-220 nm in length, called protofibrils (1,5,17).

While synthetic Aβ_1-42_ peptides are commonly used to induce neuronal toxicity in various neuronal culture systems, such as porcine brain capillary endothelial cells and co-cultures of human-induced pluripotent stem cell (hiPSC)-derived primitive macrophages with hiPSC-derived cortical neurons or cortical organoids (23,57,81), these 2D models often fail to accurately represent the native composition of neurons and glial cells. Traditional models for studying AD pathology primarily rely on *in vivo* genetically engineered mouse models (12,66,87) or the *ex vivo* 3D OBSC derived from these models (16,54,55). Some studies have explored the effects of Aβ peptides on OBSC from non-disease model mice (21,22) however, mouse models have inherent limitations as they do not fully recapitulate the metabolic pathways associated with AD pathology in humans (15,36,91). So, we have for the first time established OBSC from hippocampus region of pig brains that are much closer to humans in anatomy and physiology (79). Pigs also possess a metabolic pathway that can lead to the development of AD (23–25) and also supported our findings in OBSC developed form human hippocampus region.

Although the entry of iron into the brain is tightly controlled at the blood-brain barrier (29), the iron concentration in the brains of individuals of AD is increased 1.5 to 2-fold (3). The excess iron is localized at Aβ plaques as was confirmed using a variety of techniques using high-end technologies like electron microscopy, X-ray magnetic circular dichroism, and X-ray microscopy (2,20,64,83). Our current understanding of the contribution of the excess iron to the progression of AD is limited: on the one hand, high levels of iron are known to have an influence on both the expression of APP mRNA (33) and the activity of the α, β and γ-secretases’ (48) cleavage of APP (35–38), leading to the presence to an excess of Aβ. On the other hand, iron can bind to the hydrophilic N-terminal region of the Aβ peptide and promote the formation of oligomers and fibrils by making the peptides chemically sticky (8,42). However, little is known of the effect of iron overload on the toxicity of Aβ on the different cell types present in living and functioning nervous tissue.

To study this effect, we optimized a novel organotypic brain slice culture (OBSC) as a 3-dimensional *ex vivo* platform in which both neurons and glial cells are preserved in its native connectivity and cellular organization thus including all the cell types that normally make up the tissue in their normal arrangement. Given that the hippocampus plays a pivotal role in AD (69), we cultivated the brain slices from the hippocampus region of porcine brains and validated our findings within the same region from post-mortem human brains. Pigs demonstrate a remarkable similarity to the human brain in terms of genetics, brain morphology, size, and function (26,42).

To understand the fate of the Aβ under the impact of excess iron, we induced artificial conditions of AD pathology using by inducing synthetic Aβ_1-42_ peptides to imitate the gold standard biomarker of the disease, in addition excess iron conditions were mimicked using ferric citrate as the iron donor. Ferric citrate provides iron in ferric form which is less toxic than the ferrous form to minimize toxicity due to iron donation per se.

Using a combination of specialized light and electron microscopic techniques and cellular assays on OBSC, we demonstrate significant differences in cytotoxicity caused by Aβ_1-42_ and iron between neurons and glial cells; furthermore, we confirm in vitro that iron significantly promotes Aβ_1-42_ polymerization. For the first time, we described in the human hippocampal slices that the neurons but not glial cells internalized the Aβ. We demonstrate in vivo that the combination of iron and Aβ_1-42_ significantly increases toxicity, directing it towards neurons.

While many studies employ 2D cell cultures or mice models to primarily focus on Aβ-induced neuronal loss without taking into consideration the role of iron, our investigation sheds light on the synergy between excess iron and Aβ in the 3D *ex vivo* OBSC models mimicking AD characteristics.

## Materials and methods

### Human brain samples

The whole human brains were acquired through two routine brain autopsies conducted at the Diagnostic and Research Institute of Pathology, University Hospital Graz, Austria. These individuals had no history of neurological disorders. The age, sex and cause of death of the deceased individuals were not considered factors for this study. The brain samples were only included if the post-mortem interval (PMI) was less than 6 h at the time of collection. Additionally, it took 4 to 5 h from the time of specimen collection from pathology to the time of cultivation. Hence, the PMI here was considered to range anywhere between 5 to 15 h. In each case, the hippocampus region was carefully identified and dissected with the help of pathologists and transferred immediately to a falcon tube filled with ice-cold 0.9% saline solution buffer. This isolated region was chosen for subsequent analyses using the OBSC technique. This study was approved by the Ethics Committee of the Medical University of Graz, Austria, under votum number 28-549 ex 15/16 and 31-558 ex 18/19 and was performed in accordance with the ethical standards as laid down in the 1964 Declaration of Helsinki and its later amendments or comparable ethical standards.

### Porcine brain samples

The brain samples were sourced from 3 pigs undergoing standard meat sacrification procedures at a local slaughterhouse in Graz, Austria. The whole brains, collected in two halves are transported to the laboratory in 0.9% saline solution on ice. The hippocampus region was carefully identified, dissected, and was chosen for subsequent analyses using the OBSC technique. Since the brain specimens obtained from pigs were collected from a local animal slaughterhouse, where the brain is typically discarded as an unused product, no special permit was necessary.

### Organotypic brain slice culture

The isolated hippocampus region of the brain either from humans or pigs, was quickly rinsed in 0.013 M, pH 7.4 Phosphate-buffered saline (PBS) solution. The isolated hippocampus region of approximate size 1×1×1 cm^3^ was put into a freshly prepared 10% porcine skin gelatin (Sigma Aldrich, G6144-100G) prepared in Neurobasal^TM-A^ medium (Thermo Fischer Scientific, 10888022). This ensures precise cutting on Vibratome (Leica, VT1200S). After letting it set on ice, the gelatin mould was securely fixed to the stage using cyanoacrylate glue, and the stage holder was filled with 0.013M PBS. A stainless-steel razor blade was used as a knife. For optimal sectioning, the vibration amplitude was set to 0.2 mm. Once the tissue was positioned parallel to the blade, sections with a final thickness of 150 µm and a diameter of 3-4 mm were carefully obtained. 3-4 sections were promptly mounted onto each Millicell-cell culture insert (30 mm, 0.4 µm Millicell, Sigma-Aldrich) within a 6-well cell culture plate. The culture medium used consisted of Neurobasal medium, supplemented with 2% B27, 0.25% Glutamax, and 0.2% Neuromycin, 1 ml per well. The medium was initially replaced the following day and subsequently every alternate day throughout the course of the cultivation. 83 µM ferric citrate (Sigma-Aldrich, F3388) prepared in the culture medium was used to induce iron-overloaded condition in the brain slices. Induction of Aβ deposits was done with Aβ_1-42_, which is the most toxic among its isoforms and closely associated with AD pathology (51,62,75). 35 µM synthetic Aβ peptides (Hoelzel-biotech, RPE-A-1163-2) were used to induce Aβ fibril formation in the brain slices. These peptides were reconstituted in a solution of 50 mM Tris and 150 mM NaCl, with a pH of 7.4, before subsequently mixing in the culture medium. Study consisted of 4 groups – Untreated control, Aβ, iron and a combination of both Aβ and iron. Treatments with either excess iron, Aβ or both were added to the medium beneath without disturbing the inserts (Fig. 1). Cultures were incubated at a temperature of 37°C with 5% CO_2_ for 5 days, with harvesting conducted on the 6th day for subsequent experiments. To prepare the brain slices for harvesting, the membranes were flooded with approximately 1 ml of either Neurobasal medium or PBS. Consequently, a perforated spoon (Fine Science Tools, 10370-19) was used to carefully detach the slices from the membranes as a whole and transfer them into the required container or medium for further experiments.

**Fig. 1.**
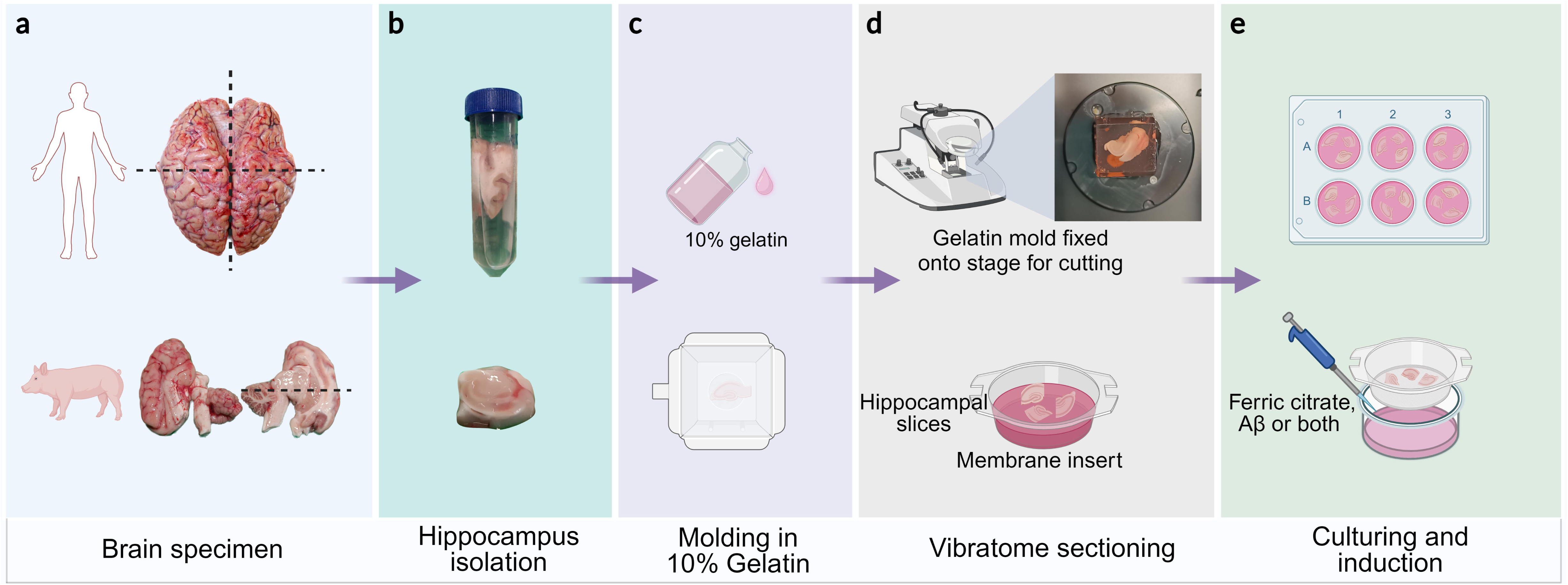
Workflow of organotypic brain slice culture (a) Obtaining the sample (b) Isolating the hippocampus region (c) Moulding into 10% gelatin (d) Vibratome sectioning at 150 µm thickness (e) Transfer to a 6-well plate and incubation for two days; ferric citrate and/or Aβ treatments via the cell culture media

### Transmission Electron Microscopy

Brain slices were fixed in a chemical fixative solution containing 2% paraformaldehyde and 2% glutaraldehyde in a 0.1 M sodium cacodylate buffer for 2 h at room temperature (RT). Following the fixation process, the samples were rinsed with 0.1 M sodium cacodylate buffer for 2 h at RT. Alternatively, the samples were kept in the rising buffer at 4°C until processed further. Samples were subsequently post-fixed using 1% osmium tetroxide solution within the same buffer for an hour at RT. To facilitate the embedding process, the samples were dehydrated using a graded series of alcohol, immersed in propylene oxide, and embedded in TAAB embedding resin (TAAB, Aldermaston, UK), all three steps were processed at RT. After allowing the resin to cure for three days at a temperature of 60°C, semi-thin sections of thickness 250 µm were carefully cut using either a Leica UC6 or a Leica UC7 ultramicrotome. The semi-thin sections were stained with toluidine blue for light microscopic analysis before proceeding with ultra-thin cutting of sections with a thickness of either 60 or 70 nm. To enhance contrast, the sections were treated with platinum blue and lead citrate solutions.

For High-Pressure Freezing and Freeze Substitution (HPF-FS), a 3 mm biopsy punch (Kai Medical) was used to make neat sections of 3 mm diameter from the harvested brain slices. These sections were sandwiched in 3 mm aluminium carriers (Engineering Office M. Wohlwend, Sennwald, Switzerland) using hexadecene as a filler to avoid air bubble formation. The sandwich was shot into the freezing chamber of an HPM 100 high-pressure device (Leica). The pressure was maintained at 2000 bar and the temperature at -196°C. After freezing the sections at high pressure, the specimen, now inside the aluminium sandwich was processed for freeze substitution (FS) with an AFS2 (Leica, Vienna, Austria) that had an inbuilt agitation device. The sample-holding agitation chamber was filled with ethanol as a conducting medium. In brief, the samples were slowly brought from -196°C to -86°C for 26 h first in a mixture of acetone and 0.1% tannic acid. This was followed by thawing the sample further to RT with an initial increase of 30°C per hour in a cocktail of acetone, 2% osmium tetroxide, and 0.2% uranyl acetate. The samples were washed in acetone and immersed in TAAB embedding resin at increasing concentrations from 50% to 100% pure resin. Then, the resin was cured for 3 days at 60°C.

For Negative staining, ferric citrate (83 µM) was mixed with the pre-prepared Aβ (35 µM) solution in the neurobasal medium. This mixture was incubated for 24 h at 37°C & 5% CO_2_. Controls included the incubations of the same preparations but at 4°C. Next, a 5 µl solution of the prepared mixture was carefully pipetted onto glow-discharged copper 400 mesh grids and allowed to incubate at RT for a minute. Excess liquid was removed by blotting with Whatman’s filter paper and stained with 5 µl of 1% uranyl acetate.

Electron microscopy was performed with either a Zeiss EM 900 at 80kV or a FEI Tecnai G2 20 electron microscope (Thermo Fisher Scientific, USA) operated at 120 kV, using BM-ultra Camera (Ultrascan 1000, Gatan, Pleasanton, USA).

### Live/dead Viability/Cytotoxicity Assay

Cytotoxicity of iron overload and Aβ fibrils in our hippocampal brain slices was assessed using InvitrogenTM’s LIVE/DEAD™ Viability/Cytotoxicity Kit (Thermo Fischer Scientific, L3224). This kit utilises chemical dyes, calcein-AM, and ethidium homodimer, to stain viable cells in green and dead cells in red, respectively. The slices were detached from the membranes as a whole and transferred, two slices per condition, into each well of a fresh 6-well plate filled with 1 ml of D-PBS. The slices were then briefly washed again with D-PBS. To prepare the staining solution, 0.5 µl of calcein and 2 µl of ethidium homodimer were added per 1 ml of D-PBS. 1 ml of staining solution was used per well to cover the entire tissue and incubated at RT for 35-45 minutes. Post-incubation, the slices were flooded with 1 ml of D-PBS to remove the staining solution. Mounting glass slides were prepared using self-adhesive reinforcement rings (Fig. 2). To prevent the compression of brain slices, the tissue slices were carefully placed inside the rings and covered with a cover slip (thickness 0.17 mm) using ProLong™ Gold Antifade Mountant from Thermo Fischer Scientific (P36934). The slides were scanned the same day using a confocal laser scanning microscope (Olympus FV3000).

**Fig. 2.**
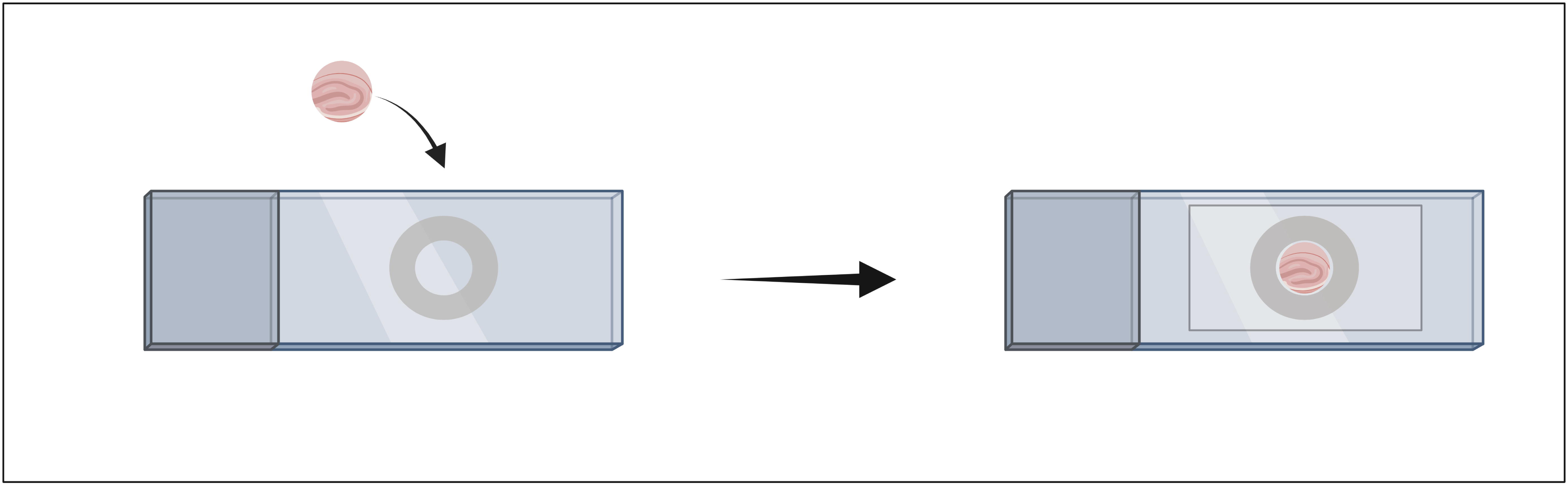
Glass slide prepared for mounting brain slices for live/dead cell viability/cytotoxicity assay

### Lactase Dehydrogenase Assay

The evaluation of lactate dehydrogenase (LDH) activity in the hippocampal brain slices under different treatment conditions was performed using the lactate dehydrogenase activity assay kit (Sigma-Aldrich, MAK066). To prepare the samples for the assay, approximately 20-30 mg of the hippocampal brain slice was homogenised in 200 µl of lysis buffer using pre-cooled beads (Qiagen, Stainless steel beads, 5 mm). Subsequently, the homogenate was transferred to a fresh tube and centrifuged at 4°C and 13,000 rcf for 10 minutes. The resulting supernatant, containing the cellular lysate, was collected into fresh tubes for further analysis. The LDH assay reaction was set up using assay components and reaction mixture as per the manufacturer’s instructions. After the incubation period, the absorbance of the samples was measured at 450 nm using a spectrophotometer.

### Immunofluorescence labelling

Immunofluorescence labelling was conducted on the hippocampal brain slices to specifically label Aβ and neurons. The protocol used for this procedure was partially adapted from (24). The brain slices were fixed by replacing the cell culture medium beneath the membrane inserts with 1 ml of 10% formalin solution and incubated at RT for 30 minutes. Following this, the formalin was completely removed, and the tissue slices were gently washed with 1 ml of 0.1 M phosphate buffer without saline at RT. The fixed tissue slices were transferred into a 24-well plate pre-filled with permeabilization solution (0.5% Triton X-100 in Phosphate buffer), with one slice per well. The 24-well plate was placed in a humid chamber and incubated overnight at 4°C. The next day, the permeabilization solution was replaced with a blocking solution consisting of 5% normal serum in PBG (0.1 M glycine, 0.8% bovine serum albumin in 0.013M PBS, pH 7.4) and 0.1% Triton X-100, and incubated overnight at 4°C. Subsequently, the blocking solution was carefully removed, and the slices were exposed to primary antibodies prepared in PBG and 0.1% Triton X-100. To label Aβ, purified anti-β-Amyloid, 1-16 Antibody (Hoelzel-biotech, 6E10) was used at a dilution of 1:200. To label the neurons, NeuN Polyclonal Antibody at a dilution of 1:200 (Thermo Fischer Scientific, PA5-78499) was used. The tissue slices were incubated in the primary antibodies overnight at 4°C.

Next, the slices were rinsed in 0.1 M PBS and incubated with secondary antibodies prepared in PBG and 0.1% Triton X-100 at RT for 4 h. We employed fluorescently conjugated secondary antibodies, Alexa Fluor™ 555 (Goat anti-Mouse IgG, Thermo Fischer Scientific) and Alexa Fluor™ 488 (Goat anti-rabbit IgG, Thermo Fischer Scientific), both at a dilution of 1:250. A subsequent washing step with 0.1 M PBS was performed to remove any excess secondary antibodies. To visualise the cell nuclei, the slices were then incubated with DAPI (Thermo Fischer Scientific) at a dilution of 1:1000 for an hour at RT and later washed with 0.1 M PBS. The tissue slices were carefully mounted onto glass slides and covered with a cover slip (0.17 mm thickness) with ProLong™ Gold Antifade Mountant (Thermo Fischer Scientific, P36934). Finally, the slides were stored in a dark environment at 4°C until further analysis. Images were obtained using a confocal laser scanning microscope (Olympus FV3000) equipped with FV31S-SW viewer software (version 2.6.1). To ensure comprehensive analysis, Z-stack imaging was conducted to gather information throughout the entire thickness of 150 µm brain slices, at wavelengths 405, 488 and 555. The resulting Z-stack layers were projected to maximum intensity images using Fiji ImageJ (Version Java 8).

### Prediction using a multi-linear regression machine learning algorithm

To assess the comparative roles of iron and Aβ in toxicity across various concentrations, we employed a multi-linear regression machine learning model. Initially, for data augmentation, we utilized the actual data, including the total cell count, and a range of ferric citrate concentrations (from 40 to 119 µM) and Aβ concentrations (from 15 to 64 µM). We trained the algorithm using this data to generate predictions for all conceivable combinations, particularly focusing on scenarios involving the concurrent presence of ferric citrate and Aβ. This encompassed combination with ferric citrate concentrations at 0 µM and varying Aβ concentrations within the specified range, as well as combinations with Aβ concentrations at 0 µM and varying ferric citrate concentrations within the specified range.

A linear regression model was trained to predict the percentage total number of dead cells, and number of neurons, using the multi-linear regression -

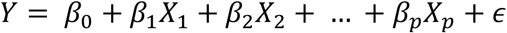

where:

Y represents the dependent variable. β_0_ is the intercept (y-intercept).

(β_1_, β_2_, ……., β_p_) are the coefficients of the independent variables.

(X_1_, X_2_, ……., X_p_) are the independent variables.

(ϵ) represents the error term.

Following this, we created a new dataset from the predicted values and attached this to the original data train. The columns used for statistical analysis were updated with the re-calculated and predicted values of the percentages of dead cells and neurons. A Gaussian Kernel Density Estimate was used to estimate the probability density function of the number of dead cells or neurons against the concentrations of either ferric citrate or Aβ for Fig. 8a–c. Linear regression analysis was performed to analyse the number of neurons to dead cells against a range of concentrations of either ferric citrate, Aβ or both in Fig. 8d.

### Statistics

Statistical analysis was performed using GraphPad Prism 9.2.0. One-way ANOVA with Tukey’s multiple comparisons test and linear regression analysis was used for Fig. 6 and 7. CellProfiler (version 4.2.5) was used to quantify the neurons and glial cells in Fig. 7.

## Results

### Intact hippocampus cellular map of post-mortem organotypic brain slice culture

To validate the usefulness of the *ex vivo* brain slice culture model for the study of neurodegenerative diseases, we examined the morphology, cellular architecture and viability of neurons and synaptic junctions in the brain slices sourced from both pigs and humans. These initial observations were crucial as impaired neuronal communication is a significant aspect of AD (31). Prior to embedding, the hippocampal brain slices from human specimens had either undergone fixation using standard chemical fixation or HPF-FS. HPF-FS was employed to mitigate any potential artefacts resulting from chemical fixation, which can lead to structural condensation due to dehydration at RT (40). HPF-FS involves rapid freezing and dehydration at low temperatures, minimizing condensation and volume changes.

In spite of the post-mortem interval, initial light microscopic observations of ultra-thin sections of human hippocampal brain slices revealed the presence of fairly dense morphology with scattered cell bodies, marked with yellow arrowheads (Fig. 3A–C). Upon closer examination of the dense regions using the electron microscope, we noted a similar dense architecture and cellular arrangement within the samples. The synaptic structures were well-preserved, with axon terminals of presynaptic neurons containing multiple mitochondria and synaptic vesicles, and evident electron density in post-synaptic neurons indicating that the brain slices maintained active synaptic communication until the fixation point in brain slices fixed via chemical fixation (Fig. 3D) and HPF-FS (Fig. 3E).

**Fig. 3.**
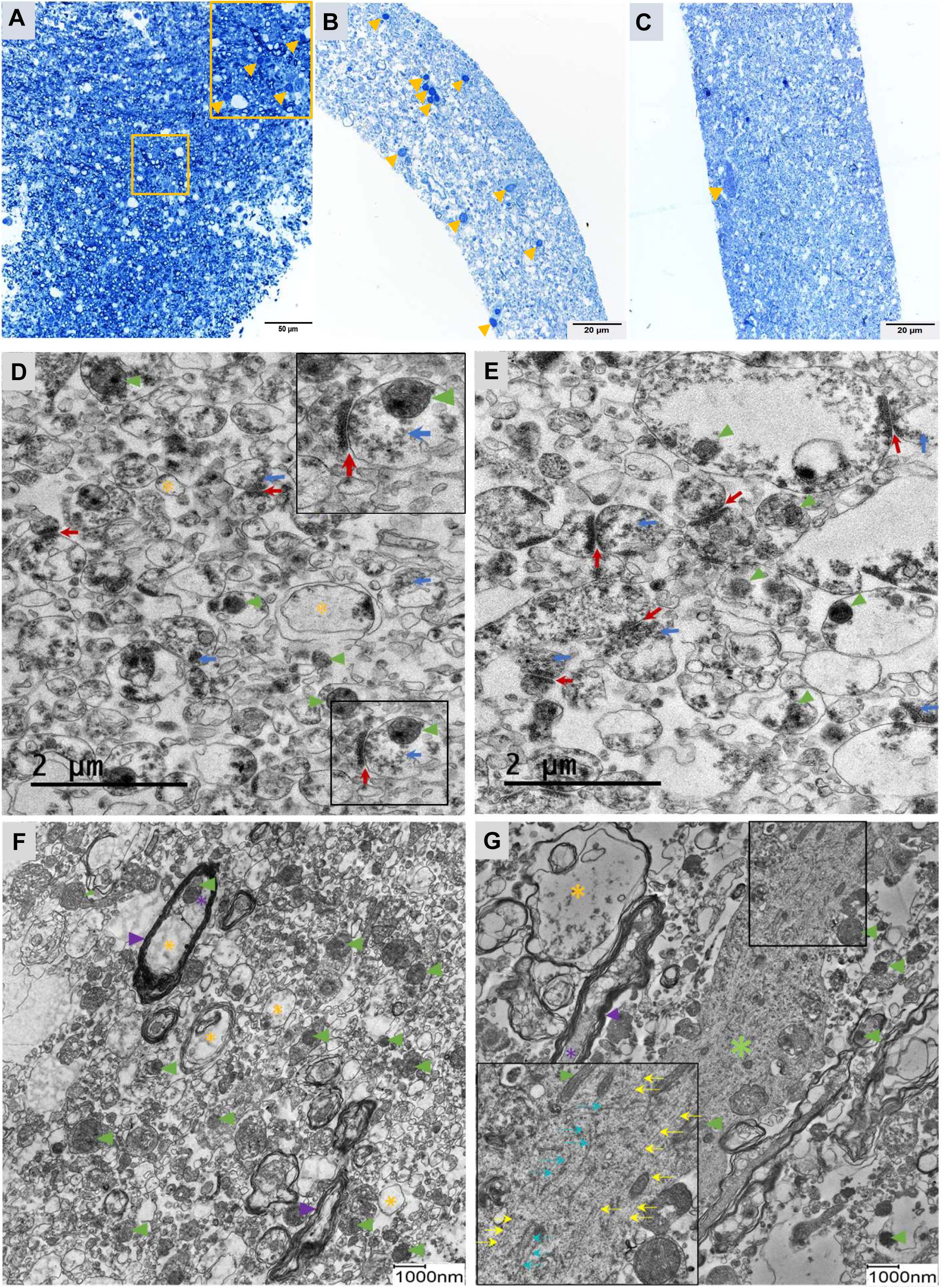
Preservation of cellular map, cellular connections and compositions Semi-thin sections of OBSC of post-mortem human brain stained with toluidine blue fixed at different intervals such as **A** fixed immediately after collection from pathology at 6 h PMI. Yellow arrowheads: cell bodies. **B** cultivated for five days via OBSC either chemically fixed or **C** fixed with HPF-FS. The brain slices display a dense cellular map with cell bodies (yellow arrowheads) uniformly spread throughout the tissue. Electron micrograph of OBSC from the post-mortem human brain **D** chemically fixed after five days of OBSC displays a dense cellular architecture with synaptic junctions marked (red arrows) and vesicles (blue arrows) inside axon terminals. Inset: synaptic junction. **E** Specimen fixed via HPF-FS after five days of OBSC displays synaptic junctions (marked with red arrows) and several vesicles inside axon terminals (marked with blue arrows). Mitochondria are visible (green arrowheads). Electron micrographs of chemically fixed OBSC from the pig brain displaying (**F**) a much denser cellular map with myelinated axons (purple asterisks) and myelin sheath (purple arrowheads). **G** Dense cytoplasm of a putative glial cell process with sub-cellular structures marked with a green asterisk. The inset shows the intermediary filaments as long, thin parallel lines (yellow arrows) and microtubules as hollow tubules (blue arrows). The mitochondria are marked with green arrowheads and cellular processes that are swollen and thus less densely filled with sub-cellular components are marked with yellow asterisks in **D–F**

The brain slices of pigs were more densely packed with cells than the human slices, but overall showed a similar cellular arrangement and synaptic structures (Fig. 3F, G). The tissue exhibited a relatively dense composition, showcasing numerous cellular processes, mitochondria, and myelinated axons (Fig. 3F). Furthermore, several cells displaying a dense cytoplasmic content in their cytoplasm (Fig. 3G, marked with green asterisks) were found, signifying a healthy and viable state of the OBSC brain slices. However, both in human and porcine samples, certain cell processes displayed a reduced number of sub-cellular compartments, as indicated by yellow asterisks (Fig. 3D–G). Within Fig. 3G, a putative glial cell process is depicted, featuring intermediary filaments that are elongated and run parallel to each other (marked with yellow arrows in the inset) and microtubules that appear as straight hollow tubes (marked with blue arrows in the inset).

### Excess iron promotes the aggregation of Aβ monomers into mature fibrillary networks in a temperature-dependent manner

Prior to initiating iron and Aβ manipulations within the hippocampal brain slices, the effects of the physiological temperature settings maintained routinely in the cell culture incubator on Aβ peptides, both in the presence and absence of iron were explored. After 24 h incubation at the cell culture incubator settings of 37°C with 5% CO_2_, in the absence of iron, we observed the formation of protofibrils characterized by various lengths and shapes, with a few starting to aggregate (marked with red arrowhead) (Fig. 4A). Protofibrils denote linear to curved prefibrillar assemblies of lengths less than 200 nm (88). In contrast, the presence of iron induced the peptides to undergo polymerization, generating elongated fibrils of various lengths. These fibrils intertwined to form intricate mesh-like networks, accompanied by a few protofibrils adjacent to the aggregated networks (Fig. 4B). Within Fig. 4C, the cross-β-helix configuration inherent to amyloid fibrils and their entwining around each other in the presence of iron at 37°C was evident.

**Fig. 4.**
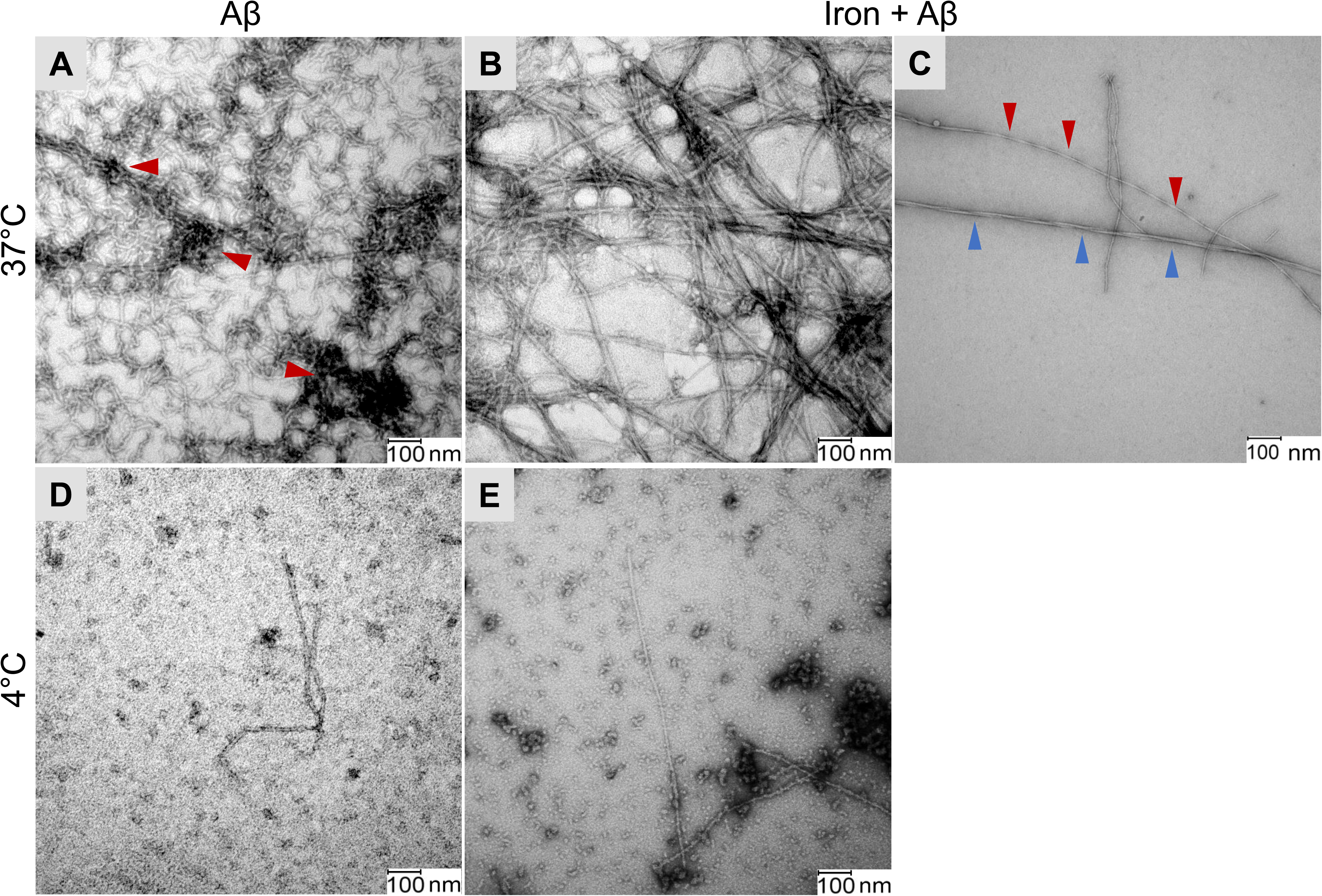
Ferric citrate promotes the aggregation of Aβ in a temperature-dependent manner Negative staining electron micrographs of **A** Aβ at 37°C in the absence of iron forming protofibrils of varying length and shape, with little protofibril aggregations (marked with red arrowheads). **B** Aβ at 37°C in the presence of iron polymerizes into a rich network of long fibrils forming thick aggregates. **C** Cross-β-helix structure of Aβ fibrils in Aβ incubated with iron at 37°C marked with red arrowheads and two fibrils intertwining with each other, marked with blue arrowheads. **D** Aβ at 4°C in the absence of iron displays an unaltered morphology. **E** Aβ at 4°C in the presence of iron forms oligomers with a few fibrils/protofibrils

At a preparatory and storage temperature of 4°C used as control, the Aβ peptides exhibited minimal changes, maintaining their native monomeric configuration (Fig. 4D) in the absence of iron. When ferric citrate was added, the peptides readily assembled into discernible globular oligomer structures, with limited formation of fibrils or protofibrils (Fig. 4E). These findings demonstrate the role of iron in promoting the formation of elongated fibrils and complex networks.

### Accumulation of Aβ fibrils in the human and porcine hippocampal region

To assess the effectiveness of Aβ treatment in a 3-dimensional hippocampal tissue culture model with a section thickness of 150 µm, the brain slices were fluorescently labelled using purified anti-β-Amyloid, 1-16 antibody to detect Aβ fibrils (Fig. 5A). The obtained z-stack images of the brain slices revealed the infiltration and distribution of Aβ fibrils across the entire thickness of both porcine samples (Fig. 5b, d) and human samples (Fig. 5f, h). We noticed that only a few cells internalized the Aβ while most of the cells did not, as indicated by the DAPI stain not surrounded by the red stain. A notable concentration of Aβ was detected at the outer edges of the tissue slice, with a comparatively lower concentration in the central optical sections. Conversely, no signals were detected in untreated control (Fig. 5a, e) and iron-treated samples (Fig. 5c, g).

**Fig. 5.**
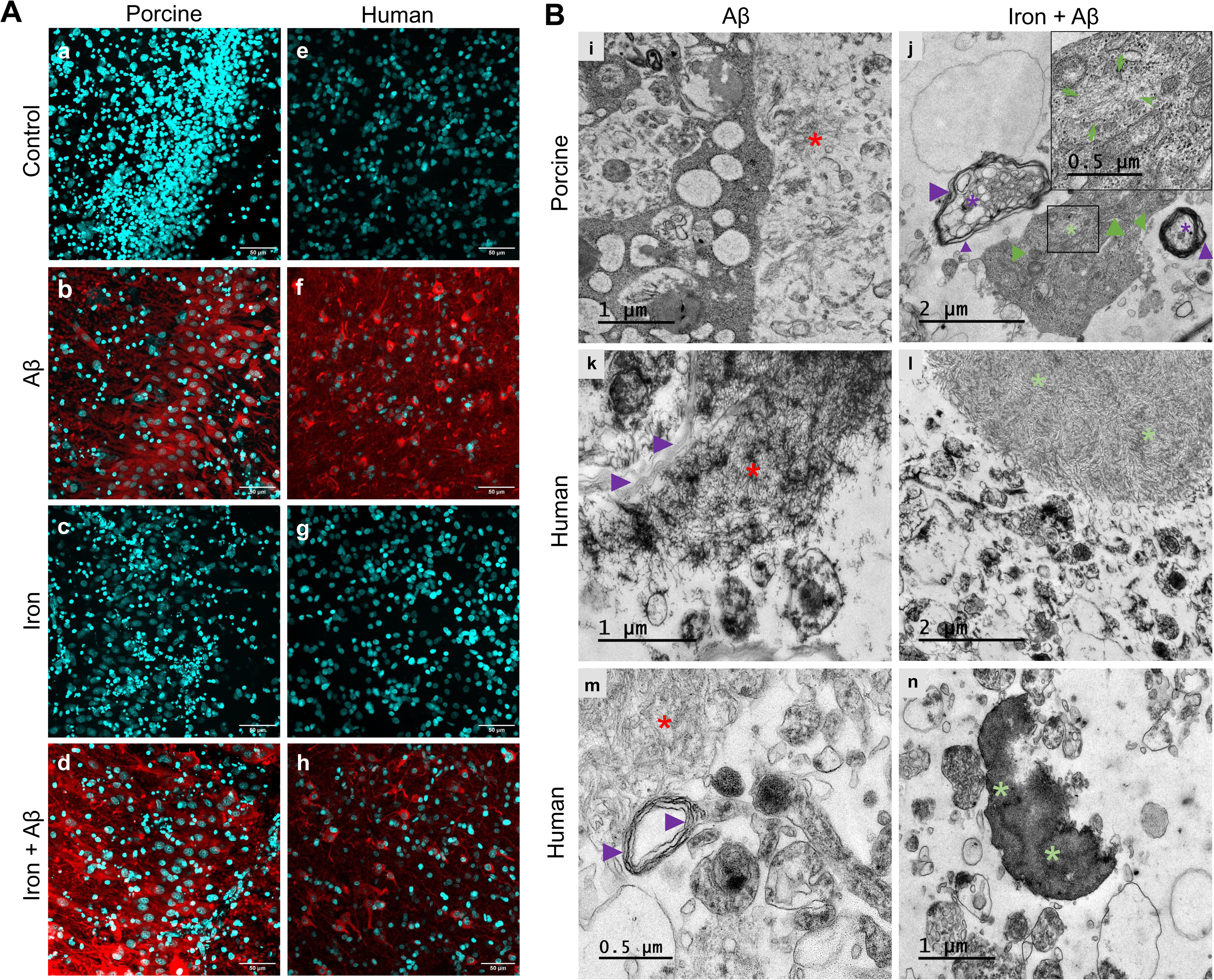
Intra- and extra-cellular accumulation of Aβ in the brain slices in human and pig brains **A** Fluorescent Aβ labelling (red) and DAPI to label nuclei (cyan) in the CA3 region of the porcine hippocampal region after treatments with (a) control (b) Aβ (c) iron (d) iron + Aβ, and in the human hippocampal region after treatments with (e) control (f) iron (g) Aβ (h) iron + Aβ. The confocal images show the intracellular accumulation of Aβ by a few cells in (b, d, f and h). Each confocal image is a maximum-intensity projection of ten z-layers with 0.55 µm z-spacing in (a) to (d) and thirteen z-layers with 0.9 µm z-spacing in (e) to (h). Scale bar: 50 µm. **B** Electron micrographs showing aggregation of Aβ fibrils in the CA3 region of the porcine hippocampal slices chemically fixed in (i) Aβ treated specimen, display disordered aggregates of fibrils marked with a red asterisk. (j) Long-rigid fibrils (green asterisks) aggregate inside the perikaryon of a neuron in the slices treated together with iron + Aβ. Aggregation of Aβ fibrils in the human hippocampus brain slices treated with Aβ (k) freeze substituted and (m) chemically fixed. The Aβ formed disordered, mesh-like aggregates (red asterisks) extracellularly surrounding distended myelin sheath (purple arrowheads), similar to the porcine specimen. Human hippocampus brain slices treated together with iron + Aβ (l) freeze substituted and (n) chemically fixed. The fibrils aggregated into a much-dense cluster within a defined area extracellularly in (l) and enclosed by a membrane in (n) marked with a green asterisk. The mitochondria inside the neuron are marked with green arrowheads. Myelinated axons (purple asterisks) are surrounded by distended layers of myelin sheath (purple arrowheads). Inset showing the long fibrillar aggregates marked within the green arrows

To obtain a closer examination of peptide polymerization and better understand if the toxicity is dependent on the chain length of Aβ, we employed electron microscopy to visualize the hippocampal slices. Our findings demonstrated that the Aβ peptides polymerized into fibrils of various lengths inside the tissue, analogous to the fibril formation observed in vitro. These fibrils demonstrated a tendency to cluster and create intricate mesh-like networks within the tissue (Fig. 5B).

In the CA3 region of the hippocampus from the porcine brain, extracellular Aβ fibrils formed disordered mesh-like structures (indicated by a red asterisk) surrounded by cellular debris and partial compartments, similar to the findings in the human specimen (Fig. 5i). In the group treated with ferric citrate + Aβ, we noted that long, rigid fibrils aggregated within the perikaryon of a neuron (Fig. 5j, indicated by green arrows). In the human hippocampus brain specimen treated solely with Aβ, the Aβ fibrils extended into long polymers and aggregated into mesh-like structures (indicated by red asterisks) extracellularly, both in specimens fixed via both HPF-FS (Fig. 5k) and through standard chemical fixation (Fig. 5m). In contrast to *in vitro*, the fibrils formed networks instead of protofibrils inside the tissue. This could be due to the presence of base levels of iron or other metals that can promote fibril formation in the tissue, e.g. see (21,26) . In the group treated with ferric citrate + Aβ, the fibrils aggregated into a much-dense cluster (indicated by green asterisks) within a defined area, either without a noticeable border as seen in the HPF-FS-fixed sample (Fig. 5l) or encased by a membrane as observed in the chemically fixed specimen (Fig. 5n). No such networks or aggregates were found in the untreated control and iron-treated samples.

### The combination of iron and Aβ is a major contributor to cell death surpassing the effects of individual treatments of iron or Aβ

The assessment of cytotoxicity for our interventions involving iron, Aβ, or their combination was performed using the live/dead cell viability and cytotoxicity assay (Fig. 6A) on porcine brain sections. The untreated control group exhibited a nearly equal number of viable and deceased cells. In the control group, the mean count of dead cells was 415 and the mean number of viable cells was 511. In the group treated with Aβ, the mean count of dead cells increased to 751 with respect to the control group while viable cells remained at 387, similar to the control group, with no statistically significant differences observed. In contrast, the iron-treated group displayed notably elevated cell death, with an average of 894 dead cells (significantly different from the control group with a p-value of 0.0076) and only 161 viable cells. The highest cell death among all conditions was detected in the combined treatment of the Iron + Aβ group, with an average of 1136 dead cells, statistically significantly different from the control group (p-value < 0.0001) and 196 viable cells, respectively. The combined treatment was significantly more toxic than the Aβ treated group, indicated by a statistically significant rise in dead cells (p-value of 0.0377) and by the presence of more viable cells in Aβ than in the combined treated group (Fig. 6B).

**Fig. 6.**
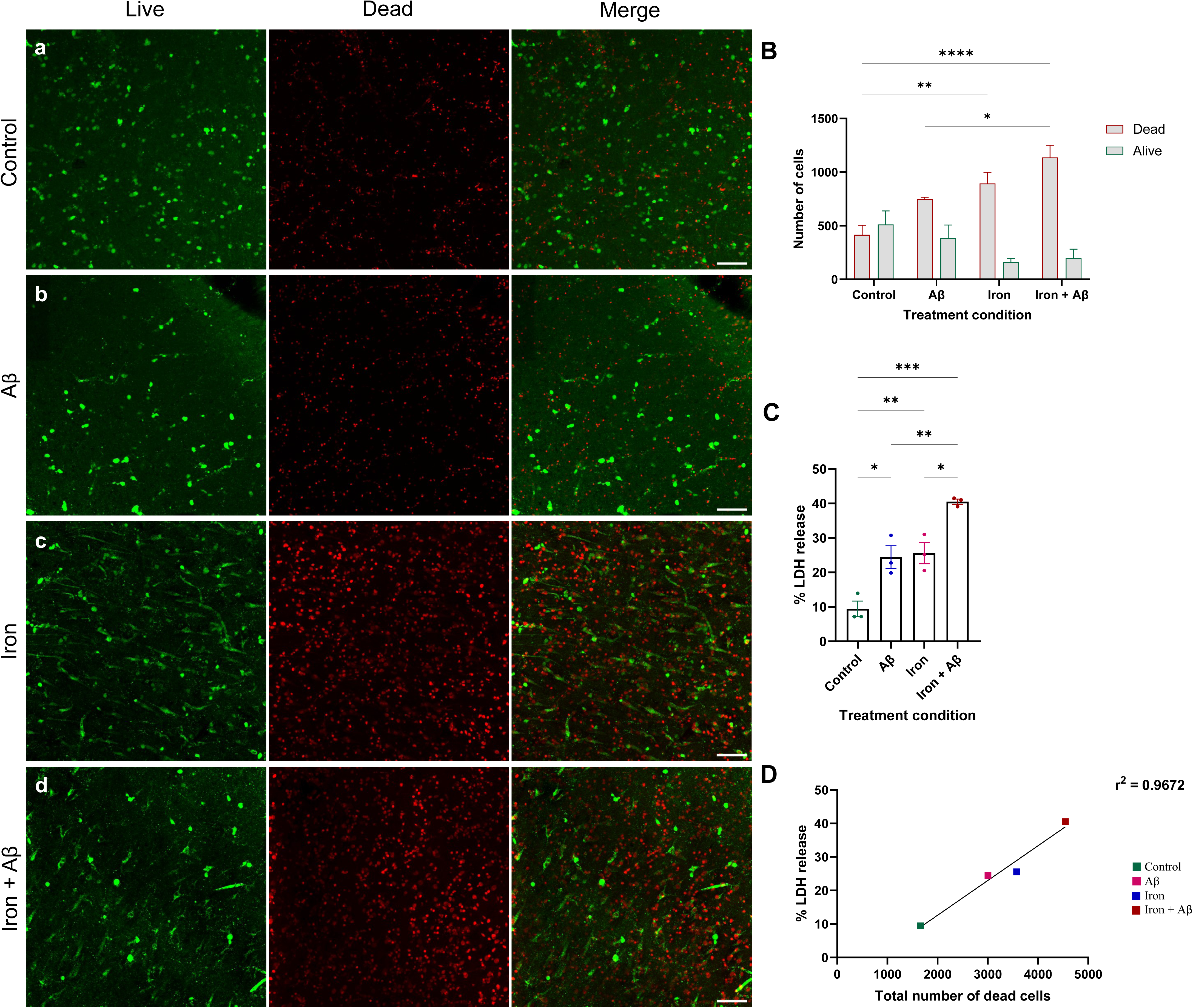
Toxicity of the combination of Aβ and iron outweighs the toxicity of their individual components **A** Confocal images of live/dead viability/cytotoxicity assay performed in porcine hippocampus brain slices treated with (a) control (b) Aβ (c) iron (d) iron + Aβ. All confocal images are maximum-intensity projections of fourteen z-layers with 2.30 µm z-spacing. Scale bar 80 µm. **B** The mean concentrations of the number of viable and dead cells plotted against the treatment condition; mean values plotted, whiskers representing SEM (n = 4). Note that the number of dead cells is highest after combined treatment followed by iron-only treatment. **C** The percentage of LDH released plotted against the treatment condition; mean values plotted, whiskers representing SEM (n = 4). In comparison to the control group, all other groups treated with either Aβ, iron, or both display a significant rise in the percentage of LDH released. The treatment groups Aβ-only and iron-only had similar percentages of LDH release while the combined treated group showed a significant increase in the percentage of LDH released when compared to iron-only or Aβ-only treated groups. (*p < 0.05; **p < 0.01; ***p < 0.001, ****p < 0.0001; One-way ANOVA with Tukey’s multiple comparisons test was used for B and C). **D** A strong positive correlation between the percentage of LDH released and the total number of dead cells confirms that as toxicity rises, so does the number of dead cells. r^2^ = 0.9672, simple linear regression analysis was used (n = 4)

To validate the cytotoxicity results, the percentage of LDH released into the ECM – indicating cytotoxicity and cell damage - by the impaired cells was analyzed. In the control group, LDH release was 9.43%. Comparatively, both the iron and Aβ treated groups exhibited higher LDH release, amounting to 25.58% and 24.45% respectively. While the LDH release in the iron and Aβ treated groups were notably similar, indicating a comparable degree of cell damage, they were significantly higher than the control group with p-values of **p < 0.01 and *p < 0.05, respectively. The highest percentage of LDH release was noticed in the fourth group, combined treatment with iron + Aβ with 40.52% of LDH release, signifying the highest level compared to the control group with a p-value of ***p < 0.01 and *p < 0.05, respectively (Fig. 6C). To establish a comparison between cell death and cytotoxicity, the data obtained through the two distinct methodologies were plotted against each other. As expected, the total sum of dead cells increased proportionately with the percentage of LDH release, with an r^2^ value of 0.9672 (Fig. 6D).

Due to the limited availability of human hippocampal brain specimens, we could only conduct the live-dead assay on human tissue once. Nevertheless, this single observation yielded an intriguing trend similar to that observed in the porcine sample (Supplementary Fig. 1A, a–d) after five days of cultivation. When comparing the number of viable and dead cells, we noted a consistent pattern between the porcine and human samples across all treatment cases, except for Aβ. In the untreated control group, the number of dead and viable cells was 481 and 369, respectively. In the iron-treated group, the number of dead and viable cells was 556 and 116, respectively. In the iron + Aβ group, the number of dead and viable cells was 390 and 42, respectively. Interestingly, in the Aβ group, the number of dead cells was higher (1270) than in the corresponding porcine sample, while the number of viable cells was 139 (Supplementary Fig. 1B, C).

### Neurons but not glial cells internalize Aβ

Through our investigation utilizing the live/dead cell viability and cytotoxicity assay, we noticed that the number of neurons appeared to decrease in samples treated with iron-only and iron + Aβ, in comparison to the untreated control and Aβ groups. Neurons were easy to identify due to their unique shape. This prompted us to investigate the susceptibility of neurons to Aβ uptake. The brain slices were fluorescently labelled with neuronal-specific marker NeuN, to detect the neurons and purified anti-β-Amyloid, 1-16 antibody to detect Aβ. Additionally, we counterstained with DAPI to visualize all nuclei. A total volume of 0.000329 mm^3^ was analyzed per treatment group (Fig. 7).

**Fig. 7.**
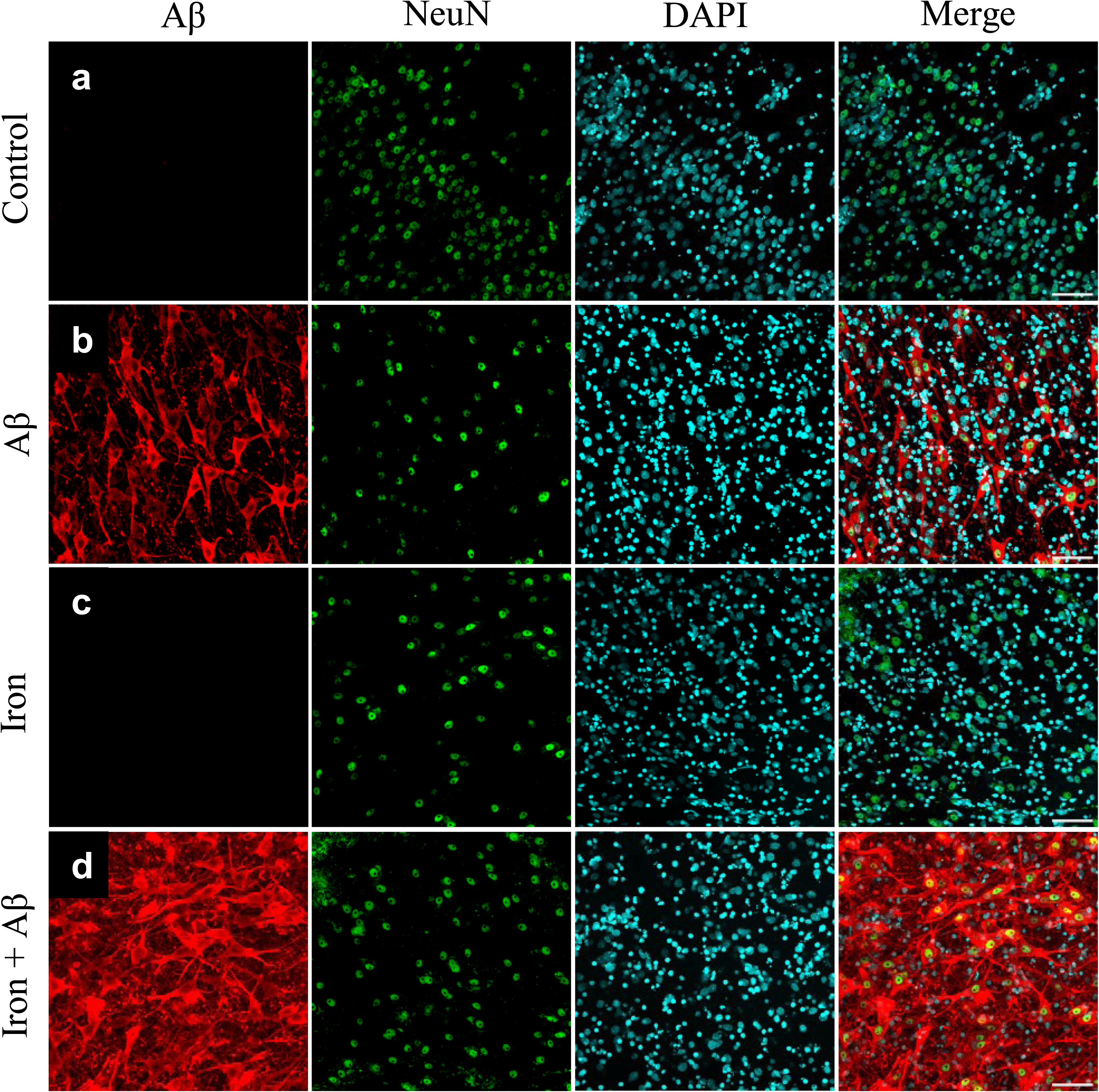
Internalization of Aβ by neurons Confocal images of double immunofluorescent staining of porcine hippocampus brain slices using an anti-Aβ antibody (red), NeuN (green), and DAPI (cyan). In the slices treated with (a) control and (c) iron, Aβ is absent but neurons are present among glial cells. Slices treated with (b) Aβ and (d) iron + Aβ show the accumulation of Aβ around the neuronal nuclei filling up the cell bodies, axons, and dendrites. Each confocal image was a maximum-intensity projection of five z-layers covering a volume of 0.000329 mm3. Scale bar 80 µm.

In the untreated control and iron groups, which had no Aβ, a red signal was absent (Fig. 7a, b). Only the neuronal nuclei stained with NeuN and all cell nuclei stained with DAPI were visible in these two groups (Fig. 7a, b). Fascinatingly, in both Aβ and iron + Aβ groups, our findings revealed that the neuronal nuclei were immediately surrounded by Aβ staining, often filling the cell body and its surrounding processes (Fig. 7c, d). Thus, in both Aβ and iron + Aβ groups, most of the interneurons and pyramidal neurons residing in the CA region of the hippocampus exhibited uptake of Aβ, while glial cells did not appear to take up Aβ fibrils. All those nuclei that do not overlap with the NeuN are composed of glial cells.

### Aβ on its own or in combination with iron selectively targets neurons

We performed a quantitative analysis of the data depicted in Fig. 7 by quantifying the mean number of glial cells and neurons present in each treatment group in a total volume of 0.00139 mm^3^ per treatment condition (Fig. 8A). The mean number of glial cells in the groups treated with Iron, Aβ, or both had a similar count compared to the control group, with the number of glial cells ranging from 435 to 514. However, there was a noticeable loss of neurons in the treated groups compared to the untreated control group (Fig. 8A). The untreated control group had a mean number of 120 neurons. Although not statistically significant, the iron-treated group had a lower mean number of 77, while the Aβ treated group had a mean number of 101 neurons. The group treated with both iron and Aβ had the highest loss of neurons with a mean number of only 37 neurons, differing statistically significantly from the control group (Fig. 8A).

**Fig. 8.**
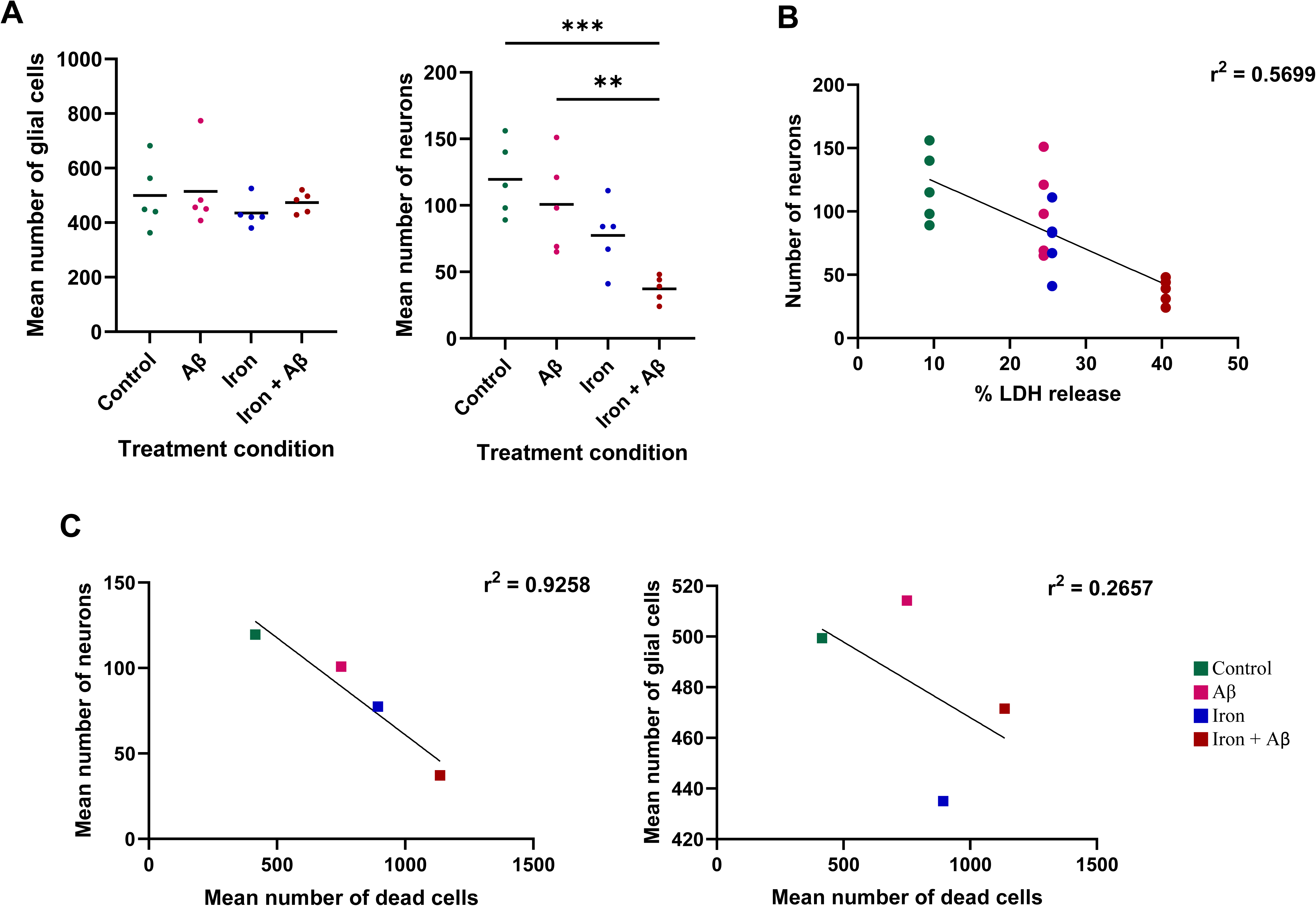
The selective impact of Aβ and iron on neuronal cells **A** The mean number of glial cells or neurons calculated from a total volume of 0.00139 mm^3^ per treatment condition (from Fig.7) were plotted against the treatment condition (Data represents mean ± SEM, n = 5). In comparison to the control group, the other three treatment conditions displayed a similar mean number of glial cells while in the case of neurons, the loss was higher in the group treated with iron than Aβ. The combined treatment group had the highest neuronal loss differing significantly from the control and Aβ treated groups. (**p = 0.0083; ***p = 0.0009; One-way ANOVA with Tukey’s multiple comparisons test). **B** As the per cent LDH release increases, there is a corresponding decrease in the number of neurons, indicating a clear association between increased LDH release and neuronal toxicity. **C** The mean number of neurons but not glial cells (n = 5) displays a strong correlation with the mean number of dead cells (n = 4). The neuronal loss is highest after combined treatment, while the glial cell loss is highest after iron-only treatment. The Aβ treated group selectively targets only the neurons but not glial cells. The r^2^ = 0.5699 in B, 0.9258 in C, left panel and 0.2657 in C, right panel, simple linear regression analysis was used

For each treatment group, the number of neurons was plotted against the percentage of LDH enzyme release to emphasize the influence of cytotoxicity on neurons. A significant correlation was observed between the percentage of LDH release, which indicates cytotoxicity, and the loss of neurons, with an r^2^ value of 0.5699. With an increase in per cent LDH release, the number of neurons decreases with the highest number of neurons in the untreated control group and the lowest in the group combined treated with iron and Aβ (Fig. 8B). To understand the relationship between neuronal loss and overall cell viability, we plotted the mean numbers of neurons and glial cells against the dead cells. The mean number of neurons exhibited an inversely proportional relationship with the mean number of dead cells, with a high r^2^ value of 0.9258. The highest neuronal death was evident after combined treatment with iron + Aβ followed by iron-only and least after Aβ-only treatment (Fig. 8C, left panel). In contrast, no correlation was observed between the means of number of glial cells and dead cells, as indicated by a lower r^2^ value of 0.2657 (Fig. 8C, right panel). The highest glial cell death was evident after iron-only treatment followed by combined with iron + Aβ and the least after Aβ-only treatment (Fig. 8C, right panel).

Interestingly the loss of neurons was most pronounced in the iron + Aβ combined treatment group, but not in the iron-only treated group, whereas, the loss of glial cells in the iron + Aβ combined treatment group fell between that of the iron-only and Aβ-only treated groups. These observations reveal that iron induces cell death in a relatively non-specific manner, affecting both glial cells and neurons. However, when combined with Aβ, this toxicity appears to spare glial cells to a large extent while selectively causing greater harm to neurons.

### Aβ directs the selectivity of iron-induced neuronal death when combined with excess iron

To dissect the differences in toxicity of iron and Aβ towards neurons, we used our experimentally obtained data to train an algorithm that predicted data on the number of neurons and dead cells and various possible combinations of iron and Aβ. The data from the quantification of dead cells and neurons were used to predict a Gaussian density curve. The curve shows the increase in the number of dead cells with rising concentrations of iron and Aβ. Although not statistically significantly different, there appeared to be a higher number of dead cells with increasing concentrations of iron compared to Aβ (Fig. 9A, B). When plotting only the number of surviving neurons the curve showed that their count decreased with increasing Aβ concentration, almost reaching zero at its peak. On the other hand, there was only a slight decrease in neuron count with increasing iron concentration, but it never reached zero (Fig. 9C). The regression analysis shows that the notable loss of neurons relative to the dead cells is higher in rising concentrations of Aβ than that of iron, as indicated by the steep and flat curves, respectively (Fig. 9D). Furthermore, with the rise in combined treatment concentrations of iron and Aβ, the loss of neurons fell between that of the groups treated with either iron or Aβ (Fig. 9D). These findings suggest that while iron-induced cell death shows a certain preference for neurons, Aβ shows a much higher selectivity in its toxicity towards neurons, maximizing neuronal loss in the presence of iron.

**Fig. 9.**
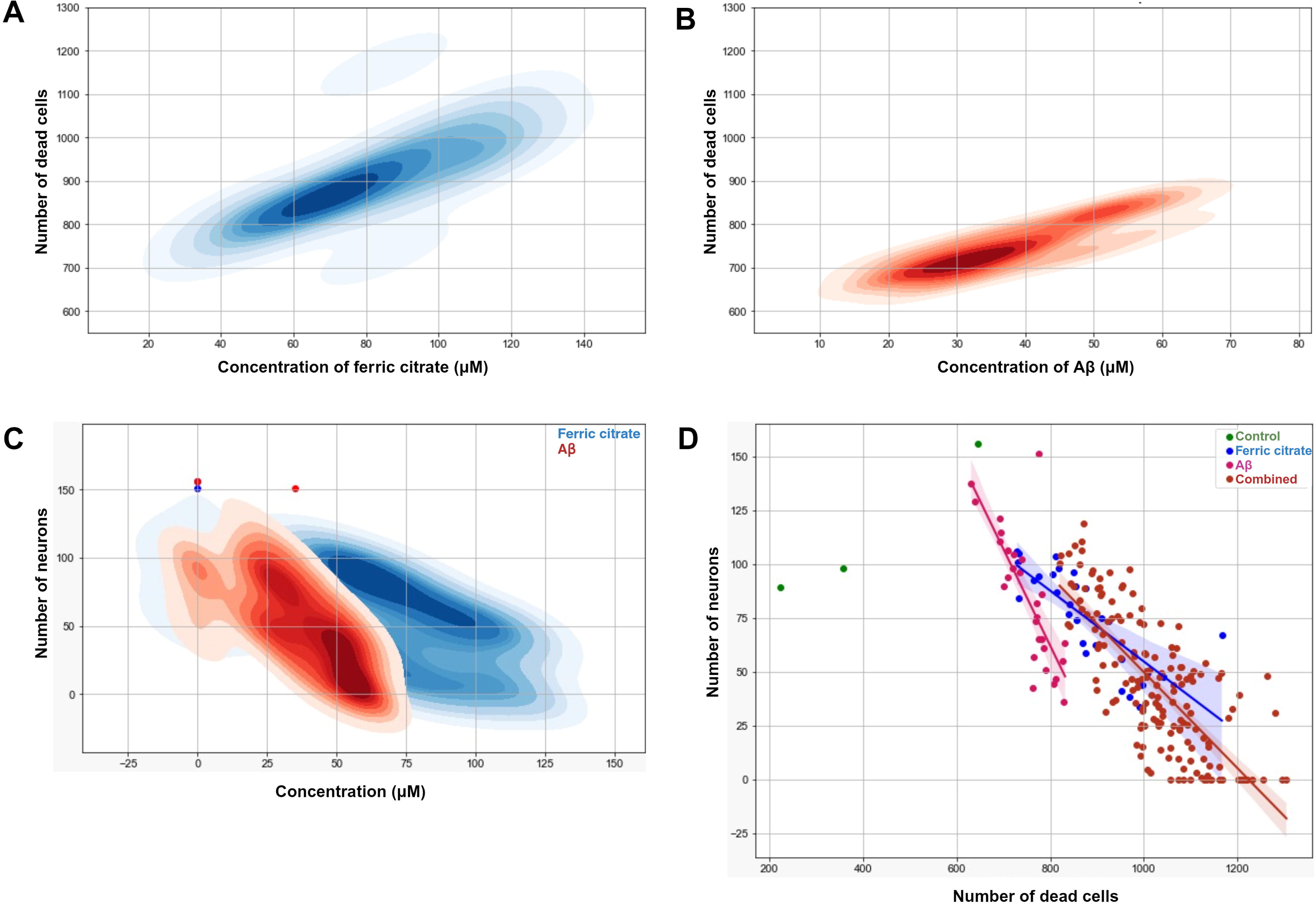
The synergistic toxic effect of iron and Aβ on neurons, with Aβ directing the toxicity Gaussian kernel density estimates from a predicted analysis of the number of dead cells plotted against the concentrations of **A** ferric citrate and **B** Aβ. The curve displays a higher number of dead cells with increasing concentrations of iron compared to Aβ. **C** The number of surviving neurons plotted against the increasing concentrations of iron or Aβ. Note that the neuronal count would reach zero with increasing Aβ concentration within the observed concentration range, whereas it decreases only slightly with increasing iron concentration, revealing that Aβ is more toxic towards neurons than iron. **D** The number of neurons plotted against the number of dead cells displays a higher loss of neurons relative to the dead cells with rising concentrations of Aβ (r^2^ = 0.6096) than that of iron (r^2^ = 0.6120), as indicated by the steep and flat curves, respectively. The loss of neurons relative to dead cells of rising concentration of combined treatment (r^2^ = 0.5893) would fall between that of the groups treated with either iron or Aβ, suggesting that the presence of Aβ along with iron directs the toxicity towards neurons. Simple linear regression analysis was used in **D**

## Discussion

The involvement of excess iron in the human brain is recognized in neurodegenerative diseases (89), but detailed information regarding the influence of iron on specific cell types in combination with Aβ remains limited. In this study, we investigated the impact of iron on neurons and glial cells and its relationship with Aβ in an artificial *ex vivo* model of AD. We demonstrated that OBSC can serve as a novel and reliable *ex vivo* model for AD drug discovery research and reported the impact of iron and Aβ on cytotoxicity, and loss of neurons and glial cells.

First, we established an OBSC suitable for investigating the role of iron and Aβ in AD. By utilizing the carefully dissected hippocampus region from the post-mortem pig and human brains, culturing brain slices via OBSC, and inducing Aβ deposits within the tissue, we were able to create an experimental model that closely reflects an important hallmark of AD, namely the Aβ deposits.

Brain explants from porcine sources proved well sustainable and reproducible. Pigs also possess a metabolic pathway that can lead to the development of AD (36), and transgenic mini pigs models have already been used as AD models (28,34,53). Moreover, they were readily available at the local slaughterhouse and provide a practical and ethical approach to obtaining brain tissue for scientific investigations.

A major advantage of OBSC over traditional mono or co culture models is that the slices retain the same cell types arranged in identical 3D configurations as found in intact tissue. We also proved that the neurons maintained intact-appearing synaptic connections even after 5 days in culture. However, in spite of proven overall viability, in porcine brain specimens, the untreated control group displayed a proportion (44.82 %) of dead cells. This could be due to the PMI causing autolysis and thus cell death, or the mechanical trauma incurred during the dissection and isolation of the region of interest within the brain tissue.

In addition to porcine samples, post-mortem human brain samples were obtained during autopsies within a 15 h PMI. They displayed the same signs of viability and connectivity as the porcine tissue, albeit with some signs of autolysis such as a swollen appearance of ECM, and distended myelin sheaths. It is important to note that autopsies are not performed immediately after death. The cadaver is usually refrigerated until autopsies. Consequently, the storage temperature partially slows down autolysis, a tissue breakdown process that starts immediately after death (14). The finding of good viability in tissue culture in spite of the PMI supports earlier studies which reported that the majority of proteins and nucleic acids remained stable up to a PMI of 30 hours, and that neurons of viable morphology remained present, and DNA repair activity remained intact up to a PMI of 24 hours (6,14,44,73,85).

To our knowledge this is the first study in which Aβ_1-42_ as well as ferric citrate was added to porcine and human OBSC in order to generate Aβ deposits and iron accumulation as a model for AD. (9,52,58,86). We obtained several lines of evidence from our study that our OBSC approach provides a good model for studying AD.

Firstly, electron microscopy demonstrated that Aβ exhibited aggregate formation within the tissue; Second, the Aβ proved toxic mimicking the neurotoxicity of AD in humans: as expected, the mean number of dead cells increased in the Aβ treated group due to the toxicity exerted by Aβ. Third, an antibody directed against Aβ proved its uptake into the tissue.

Furthermore, we found that the Aβ had been selectively taken up by neurons from the extracellular pool, but not by glial cells. To our knowledge this selective uptake into pig or human hippocampal neurons in 3D had not been described so far. The Aβ uptake by neurons could have occurred via the apoE receptors (13,25,41) or integrins and NMDA receptors in the neurons (4) as described by earlier studies conducted in the human brain. The specific form of Aβ (oligomers, protofibrils, or fibrils) taken up by neurons remained unclear as the anti-Aβ antibody binds to the 1-16 amino acids of the Aβ peptide and thus does not distinguish according to chain length. Electron microscopic visualization of Aβ in different forms, such as monomers, oligomers, or protofibrils inside the dense cytoplasmic content of neurons posed challenges, due to their resemblance in shape and size to sub-cellular components like Nissl granules and smooth endoplasmic reticulum.

The fact that Aβ was not found within glial cells was unanticipated as we expected microglial cells, the resident immune cells in the brain, or astrocytes to internalize Aβ for clearance (43,59,70).

We next combined our OBSC model of AD with an in vitro approach to demonstrate the effect of iron on Aβ structure and toxicity.

We found that iron promotes Aβ polymerization and thereby leads to fibril formation in vitro. In the OBSC experiments, fibrils were found inside neuronal cell bodies in the group treated with both iron + Aβ, and the extracellular clusters of fibrils were much denser when excess iron was present as compared to the group treated with Aβ only. These dense clusters resembled those observed in DPP6-KO mice and AD human brains (45). That iron promotes fibrillization of Aβ was also found in a proteomic study by (8) describing the formation of β-sheet structures upon binding of ferrous iron to Aβ peptides. Similarly, a study by (71) revealed comparable intertwined Aβ structure in the presence of ferric citrate using EM. In contrast, (72)showed that iron reduces chain length of Aβ, in a study that had been performed with a significantly higher ratio of iron to Aβ concentrations. Both in the presence or absence of iron, polymerization was stronger at 37°C than at 4°C, supporting existing literature on the temperature-dependence of fibril formation (73)

Of note, increased numbers of dead cells were found in the iron-treated group. This contrasts with previous studies suggesting the non-toxic nature of ferric citrate after extended oral supplementation in mice and adult rat brains (32,84). However, it needs to be born in mind that in our case, the ECM was directly exposed to the medium containing ferric citrate with no uptake control mechanisms such as the blood brain barrier. The increase in dead cells may be caused by oxidative damage induced by metal ions (65) or dysregulation of iron homeostatic proteins (33,68,77,90).

The highest rise in cell death in the OBSC model was noted when both iron and Aβ had been added together. In line with our cytotoxicity findings, the release of the LDH enzyme, a marker of cellular damage, into the extracellular matrix, increased in both the iron-only and Aβ-only treatments as compared to untreated control, and the group subjected to combined iron + Aβ treatment exhibited the highest LDH release.

### Which cell types are affected by Aβ and iron toxicity and how?

We found a similar number of glial cells but a noticeable difference in the number of neurons throughout different treated conditions. The loss of neurons was highest after iron + Aβ treatment, followed by Aβ-only and least after iron-only treatment. Click or tap here to enter text.In a novel approach making use of recent machine learning algorithms, we looked at the toxicity of Aβ, iron, and a mixture of both with predicted synthetic data. Both real and synthetic data demonstrated that excess iron on its own leads to unspecific cell death with only a slight an inclination towards neurons, whereas the presence of Aβ specifically directs toxicity towards neurons when combined with iron. Synthetic data further showed that a much lower concentration of Aβ, compared to iron, would be required to kill all neurons.

One hint as to a possible mechanism for the synergistic effect of iron and Aβ comes from our previous study that demonstrated that neurons retain more ferritin (an iron reservoir) with respect to glial cells when the total ferritin level is high (82). Thus, extra iron present inside neurons upon Aβ uptake might lead to an alteration in the structure of Aβ and thus increased intracellular toxicity. Ferritin has been shown to colocalize with senile plaques *in vitro* and in post-mortem AD brains (83,84), and *in vitro* investigations using scanning transmission X-ray microscopy and electron microscopy have provided insights into the co-aggregation of Aβ and ferritin, revealing the conversion of ferric iron in the ferritin core into more reactive ferrous ions (85,86).

### What is the mechanism by which iron promotes Aβ toxicity?

Although direct effects of iron such as oxidative stress or the dysregulation of iron-regulating proteins cannot be ruled out, the higher toxicity of a combination of iron and Aβ than of Aβ on its own can be attributed to the fact that iron causes changes in Aβ structure, thereby increasing its toxicity.

The mature Aβ assemblies formed in the presence of excess iron can interact with various receptors, such as the p75 neurotrophin receptor, metabotropic glutamate receptors, α-nicotinic acetylcholine receptor, N-methyl-D-aspartic acid receptor, and β-adrenergic receptor, (87–89) and disrupt their physiological function. A study conducted in neuronal cell cultures showed reduced toxicity upon partial reversal of Aβ aggregation (90). Fibrillization of Aβ has been shown to exacerbate synaptic communication loss (91,92), and other previous studies show that iron-induced Aβ aggregations disrupt calcium homeostasis (93), impair glutamatergic synaptic transmission, and hinder neuronal glutamate uptake, ultimately affecting synaptic plasticity and neuronal survival (81).

In summary, we present several lines of evidence indicating that iron influences Aβ polymerization and contributes to its toxicity. Firstly, *in vitro* we demonstrated that excess iron promote Aβ polymerization. Second, fibrils were more readily observed under electron microscopy when both iron and Aβ were added to the OBSC model due to their formation of dense clusters. Third, factual and predicted data give strong evidence that the combination of iron and Aβ is more toxic than either substance alone, with this toxicity being selective towards neurons. Currently, several therapeutic approaches targeting Aβ fibrillogenesis are actively pursued due to compelling evidence from *in vitro* and *in vivo* studies indicating that fibrillar Aβ can directly harm neurons (49,94). However, the role of iron in Aβ fibrillogenesis and subsequent neuronal cell death is often overlooked in drug discovery. Our findings suggest a synergistic interaction between iron and Aβ in neurons in iron-overloaded conditions, resulting in a biased toxicity towards neuronal death. The highly neurotoxic nature of iron-mediated Aβ aggregate formation underscores the need to target excess iron as a preventive measure in AD management.

Our study thus describes how iron contributes to Aβ toxicity, however more detailed studies are required to quantify the accurate contribution of iron on Aβ. Our novel, porcine and human OBSC model has the potential to serve as a platform to expand these studies further.

## Data availability

The data presented in the manuscript is fully available upon request to the authors.

## Code availability

All the scripts and software used in this study’s analysis were built upon Python and are available upon request to the authors.

## Consent to participate

Not applicable.

## Consent to publish

Not applicable.

## Acknowledgements

We thank Luca Schmid, and Nina Schlögl for expert technical assistance. We also thank Alexander Nartey for his help with confocal imaging. Fig. 1 and 2 are created with BioRender.com.

## Author contribution

Concept, project design and study plan: SS and GL. Planning and performing the experiments: SS and GL. Statistics and analysis: SS, RK, and WG. Cell culture: SS, SP. Electron microscopy: SS, WK, and GL. Human brain acquisition: ML, AB, and JH. Prepared Figures: SS, RK, SR. All authors revised, read, and approved the manuscript.

## Ethics declaration

This work was approved by the Ethics Committee of the Medical University of Graz, votum number 28-549 ex 15/16.

**Supplementary information (SI) – Fig. A1**

Confocal images of Live/dead viability/cytotoxicity assay performed in human hippocampus brain slices treated with (a) control (b) iron (c) Aβ (d) combination of iron + Aβ. All confocal images are maximum-intensity projections of seven z-layers with 3.93 µm z-spacing. Scale bar 80 µm. The number of (e) dead and (f) viable cells plotted against the treatment condition represented as a bar graph. The loss of both live and dead cells follows a similar trend to that in the pig specimen, except for the Aβ treatment group, which shows a higher number of dead cells compared to its corresponding treatment in the pig brain

