## Supplementary figures and images for "Amyloid beta aggregation promoted by iron leads to neuronal loss in an *ex vivo* model of Alzheimer’s disease"

### Supplemental Data 1

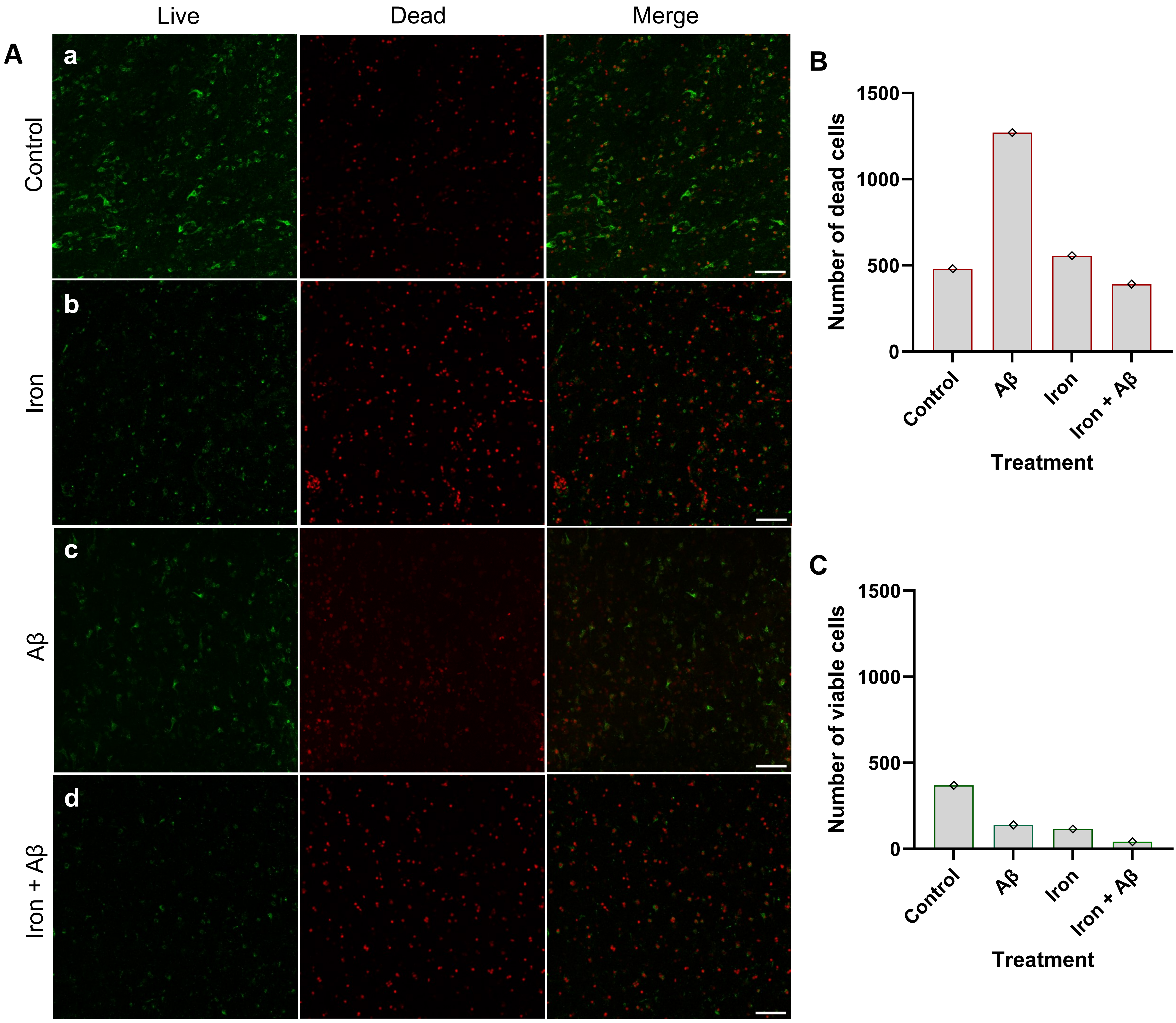
